# Making WAVES in Breedbase: An Integrated Spectral Data Storage and Analysis Pipeline for Plant Breeding Programs

**DOI:** 10.1101/2020.09.18.278549

**Authors:** Jenna Hershberger, Nicolas Morales, Christiano C. Simoes, Bryan Ellerbrock, Guillaume Bauchet, Lukas A. Mueller, Michael A. Gore

## Abstract

Visible and near-infrared (vis-NIRS) spectroscopy is a promising tool for increasing phenotyping throughput in plant breeding programs, but existing analysis software packages are not optimized for a breeding context. Additionally, commercial software options are often outside of budget constraints for some breeding and research programs. To that end, we developed an open-source R package, *waves*, for the streamlined analysis of spectral data with several cross-validation schemes to assess prediction accuracy. *Waves* is compatible with a wide range of spectrometer models and performs visualization, filtering, aggregation, cross-validation set formation, model training, and prediction functions for the association of vis-NIRS spectra with reference measurements. Furthermore, we have integrated this package into the Breedbase family of open-source databases, expanding the analysis capabilities of this growing digital ecosystem to a number of crop species. Taken together, the standalone and Breedbase versions of *waves* enhance the accessibility of tools for the analysis of spectral data during the plant breeding process.

**Core ideas:**

- *waves* is an open-source R package for spectral data analysis in plant breeding
- Breeding relevant cross-validation schemes to evaluate predictive accuracy of models
- Extension of Breedbase—an open-source database—to support spectral data storage
- Graphical user interface developed for implementation of *waves* in Breedbase

## INTRODUCTION

Visible and near-infrared spectroscopy (vis-NIRS) allows for rapid and low-cost analysis of the physical and chemical makeup of biological samples in a non-destructive manner through the calibration of prediction models relating spectra to reference values. This is especially useful in the context of plant breeding programs, which often rely on expensive and time-consuming quality trait information for the development and release of improved plant varieties. In these cases, the prediction of quality traits with vis-NIRS affords an opportunity to increase the efficiency of varietal development.

The use of NIRS to predict quality traits in agricultural products revolutionized the industry over 50 years ago (McClure 2003; Teixeira Dos Santos, Lopo, Páscoa, & Lopes, 2013), but only with the recent application of microelectromechanical systems to spectroscopy has the cost of high-quality spectrometers reached a reasonable price point for most plant breeding and research programs (Crocombe 2004; Pasquini 2018; Teixeira Dos Santos et al. 2013). In order to evaluate and appropriately deploy these new tools in research and other high-value applications, prediction models must be developed for each phenotype of interest. Though many spectrometers come equipped with proprietary models that are pre-trained for phenotypes such as total protein content or brix, low-cost hardware often requires development and maintenance of prediction models to be performed by the user. While model calibration as a service and standalone commercial software are viable solutions, the high cost of both negates the benefits of an inexpensive spectrometer.

Open-source software packages for the analysis of vis-NIR spectral data are available in several programming languages, including R (R Core Team 2020) and python (Van Rossum & Drake 2009), but thorough analysis, regardless of the language, requires functions to be systematically assembled together from many different packages or libraries. In R, a user interested in spectral analysis must pull together preprocessing functions from *prospector* (Stevens & Ramirez-Lopez 2014) and model training functions from packages such as *pls* (Mevik, Wehrens, & Liland, 2019), *randomForest* (Liaw & Wiener 2002), *kernlab* (Karatzoglou, Smola, Hornik, & Zeileis, 2004), and *caret* (Kuhn 2020). Without a working knowledge of these programming languages and training in vis-NIR reflectance analysis, the construction of a functional pipeline is unquestionably challenging. A web-based graphical user interface (GUI) would lower the entry point to spectral analysis, effectively democratizing vis-NIRS when paired with low-cost spectrometers.

We developed an R package, *waves*, that brings essential spectral analysis functions together to enable streamlined filtering, preprocessing, model training, and phenotype prediction with cross-validation schemes tailored to address prediction scenarios commonly faced by plant breeding programs (Jarquín et al. 2017). To further leverage the functionality of *waves* in a database environment without a need for familiarity with the R programming language, we also created a *waves*-based spectral data analysis tool in Breedbase, an open-source family of relational databases for the storage and analysis of plant breeding genomic and phenomic data (https://breedbase.org; see Figure 1). This integration also includes the development of spectral data storage and handling capabilities within the database, linking spectral JSONB tables with related field trial meta-and phenotypic data. The Breedbase digital ecosystem allows users to store all available data across germplasm sets and environments, which can then be accessed to customize calibration models. To our knowledge, this is the first open-source integration of spectral data storage and analysis with plant breeding database tools.

**Figure 1.**
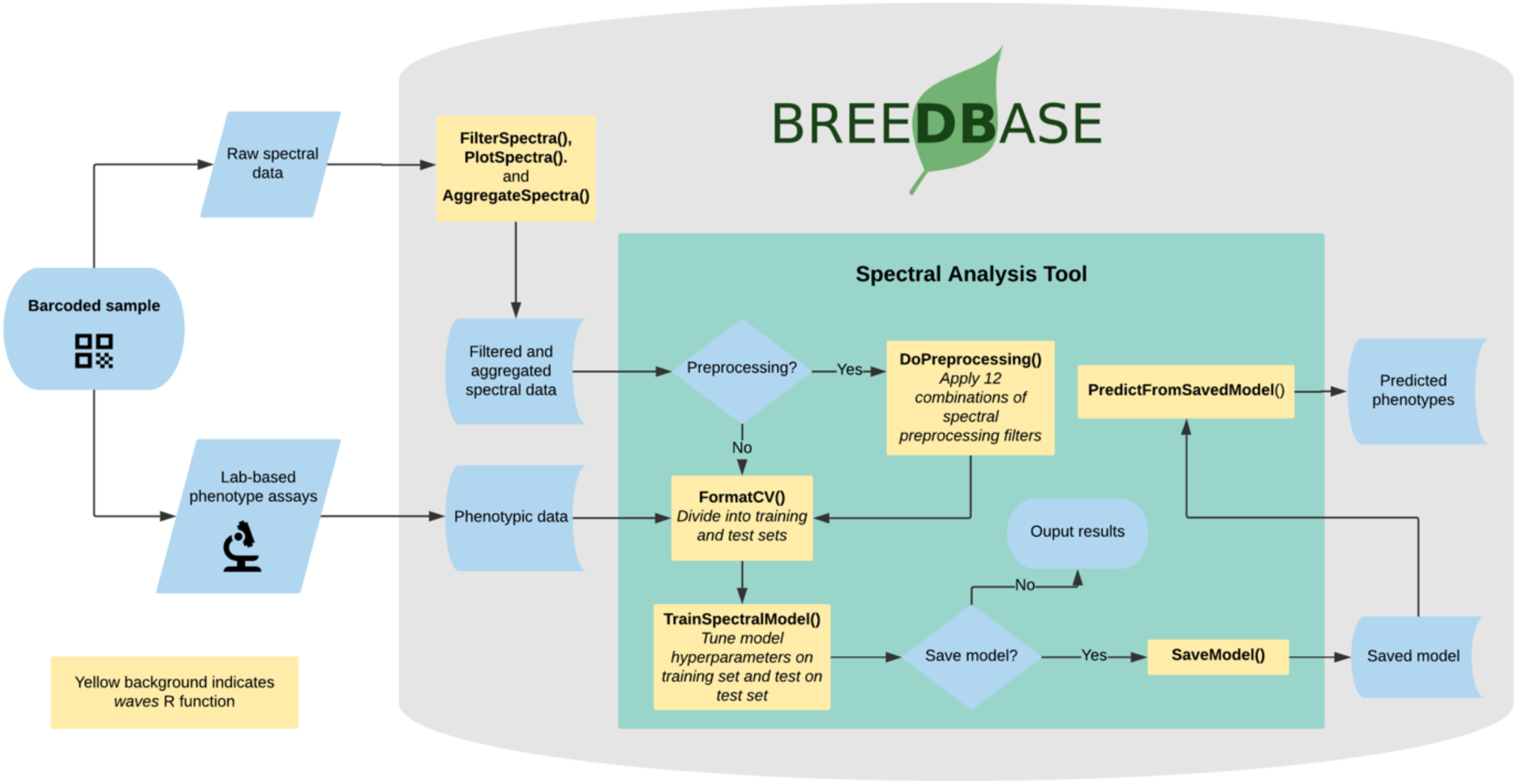
*waves* R package functions are integrated into the Breedbase spectral analysis tool for spectral data storage, preprocessing, cross-validation set formation model development, model storage, and trait prediction.

*Waves* is a standalone R package that is called by the backend Perl, JavaScript, and R code base of Breedbase (https://breedbase.org/). The user manual for Breedbase is available at https://solgenomics.github.io/sgn with information regarding the *waves* integration in the sections titled “Spectral Data Upload and Storage” and “Spectral Data Analysis,” and the user manual for the standalone R package *waves* version 0.1.0 is available in the supplemental files. The R package *waves* is available through the Comprehensive R Archive Network (https://CRAN.R-project.org/package=waves).

## IMPLEMENTATION

### Data storage

The storage of spectral data can be a major challenge for vis-NIRS users, as high-dimensional datasets are common and must be manipulated in a controlled manner to correctly execute calibration procedures. Though proprietary software often offers both storage and analysis features, open-source resources are neither linked to phenotype data nor optimized for use in analyses (Chalk 2016; https://spectralworkbench.org). Breedbase stores spectral data alongside other types of phenotypic data, eliminating the need for data matching during analysis. Breedbase instances are governed by controlled ontologies based on the Crop Ontology system (Shrestha et al. 2012). We have extended the statistical ontology for this system to include spectroscopy analysis algorithm terms (See supplementary table 1), and these terms are combined with existing trait ontology terms to generate spectroscopy-based predicted ontology terms. Spectral data can be uploaded to Breedbase as .csv files. Future data transfer using BrAPI (Selby et al. 2019) will allow for interoperability with data collection software, another step towards a complete digital ecosystem for spectral data. Given the wide variety of vis-NIR spectrometers employed by plant breeding programs, we made the system flexible enough to handle spectral data irrespective of the source spectrometer. Though our system does not handle proprietary data formats, once they are transformed into JSON or .csv, the number of wavelengths and size of gaps between them does not limit compatibility.

Spectral calibration models can be heavily affected by the presence of outliers, whether they are the result of spectrometer artifacts or user error. Mahalanobis distance (Mahalanobis 1936) is a measure of the distance between a single observation and a larger distribution and is commonly used for the objective identification of outliers in a multivariate space, including spectral datasets (De Maesschalck, Jouan-Rimbaud, & Massart, 2000). The *FilterSpectra()* function in *waves* identifies outliers by calculating the Mahalanobis distance of each observation in a given spectral matrix using the *mahalanobis()* function from the *stats* package (R Core Team 2020). In Breedbase, this procedure is applied on a per-dataset basis on upload and outliers are given binary tags “outlier.”

After outlier identification, data can be visualized using the *PlotSpectra()* function in *waves*. This function uses the filtered spectra and the *ggplot()* function from the *ggplot2* package to create a line plot with outliers highlighted by line color. A list of rows identified as outliers are shown beneath the plot. In Breedbase, these plots are saved as .png files and linked to the original input datasets. Plot image files can be downloaded via the “Download Plot” button on the upload webpage.

To obtain a stable and reliable spectral profile, most spectrometer manufacturers recommend that multiple spectral scans are captured for each sample. While some spectrometers aggregate these scans internally, many do not, requiring the user to do so before analysis can take place. Breedbase handles these cases upon data upload following filtering steps by calling the *AggregateSpectra()* function from *waves*, saving the aggregated scans for future access through the search wizard feature. After aggregation, the user exits the upload workflow and the raw data file is saved and accessible through the Breedbase file storage system.

### Analysis

In order to initiate an analysis, the user must select a dataset using the Breedbase search wizard tool. Breedbase datasets can contain observationUnit-level (plot-, plant-, or sample-level) trial metadata and phenotypic data from one or more trials. After navigating to the “Spectral Analysis” webpage under the “Analysis” tab in Breedbase, the user can select one of these datasets as input for model development.

Preprocessing, also known as pretreatment, is often used to increase the signal to noise ratio in vis-NIR datasets. The *waves* function *DoPreprocessing()* applies functions from the *stats* (R Core Team 2020) and *prospectr* (Stevens & Ramirez-Lopez 2014) packages for spectral preprocessing, giving the user the option to keep the raw data (default in Breedbase) or use any of 12 combinations of common preprocessing methods including standard normal variate (Barnes, Dhanoa, & Lister, 1989), first and second derivatives, and Savitzky-Golay polynomial smoothing (Savitzky & Golay 1964).

During analysis, the user can select from six cross-validation (CV) schemes that represent scenarios common in plant breeding using the *FormatCV()* function in *waves*. These include CV1, CV2, CV0, and CV00 as described in depth by Jarquín et *al*. (2017), as well as random and stratified random sampling. For those four schemes from Jarquín et *al*., the user must choose specific trials from their input dataset based on genotype and environment relatedness. Guides for these selections are available in the *waves* and Breedbase manuals, and they are accessible as pop-ups through the spectral analysis webpage.

Several common machine learning (ML) algorithms are available for calibration model development in Breedbase via *waves*. The *TrainSpectralModel()* function in *waves* performs hyperparameter tuning as applicable using these ML algorithms in combination with CV and train functions from the package *caret* (Kuhn 2020). The current version of *waves* supports three ML algorithms; partial least squares regression (PLSR; Wold 1982; Wold, Ruhe, Wold, & Dunn, 1984) is implemented using the *pls* package (Mevik et al. 2019), random forest regression (RF; Ho 1995) is implemented with the *randomForest* package (Liaw & Wiener 2002), and support vector machine regression (SVM; Vapnik 2000) is from the *kernlab* package (Karatzoglou et al. 2004). The modular nature of this function allows for the future inclusion of available ML approaches.

The *TestModelPerformance()* function in *waves* acts as a wrapper for the iterative testing of different selections of training and testing sets. It calls the *DoPreprocessing(), FormatCV()*, and *TrainSpectralModel()* functions and outputs model performance statistics for each iteration of set formation.

### Output

After training, common model performance statistics are both displayed on a results webpage and made available for download in .csv format. These statistics are calculated by the *TrainSpectralModel()* function in *waves* using the *spectacles* package (Roudier 2020). Once a model has been trained, it can be stored for later use. This action calls the *SaveModel()* function from *waves*. Metadata regarding the training dataset and other parameters specified by the user upon training initialization are stored alongside the model object itself in the database. Phenotype predictions can be performed by pairing saved models with spectral datasets that were collected using the same spectrometer type. Predicted phenotypes are stored as such in the database and are tagged with an ontology term specifying that they are predicted and not directly measured. Metadata regarding the model used for prediction are stored alongside the predicted value in the database. Predicted phenotypes are then available for download or use in other Breedbase analysis tools such as the Selection Index and GWAS to support decision making in the plant breeding process.

### Example dataset and performance tests

An example dataset of vis-NIR and reference phenotypic data from Ikeogu et al. (2017) is included in the package *waves*. In their study, spectra were collected from freshly sliced and shredded cassava roots using a QualitySpec Trek vis-NIR spectrometer and phenotypic reference data include root dry matter content (RDMC) using the oven method, as well as total carotenoid content (TCC) as measured by high performance liquid chromatography. When this dataset was analyzed using the *TestModelPerformance()* function of *waves* with the same training and test sets as Ikeogu et al. (2017), the predictive performance of our best PLSR models (Figure 3, Supplemental Table 2) was similar to their published best modified PLSR models as developed with commercial, proprietary software. For RDMC, the best model trained by *waves* generated an R^2^_p_ of 0.86 using a Savitzky-Golay filter, while the best model in Ikeogu et al. (2017) was generated with an SNV detrend pretreatment and had an R^2^_p_ of 0.84. Similarly, *waves* achieved an R^2^_p_ of 0.95 for TCC with a gap derivative filter, while Ikeogu et al. (2017) generated an R^2^_p_ of 0.86 using an SNV detrend filter with the same datasets.

**Figure 2.**
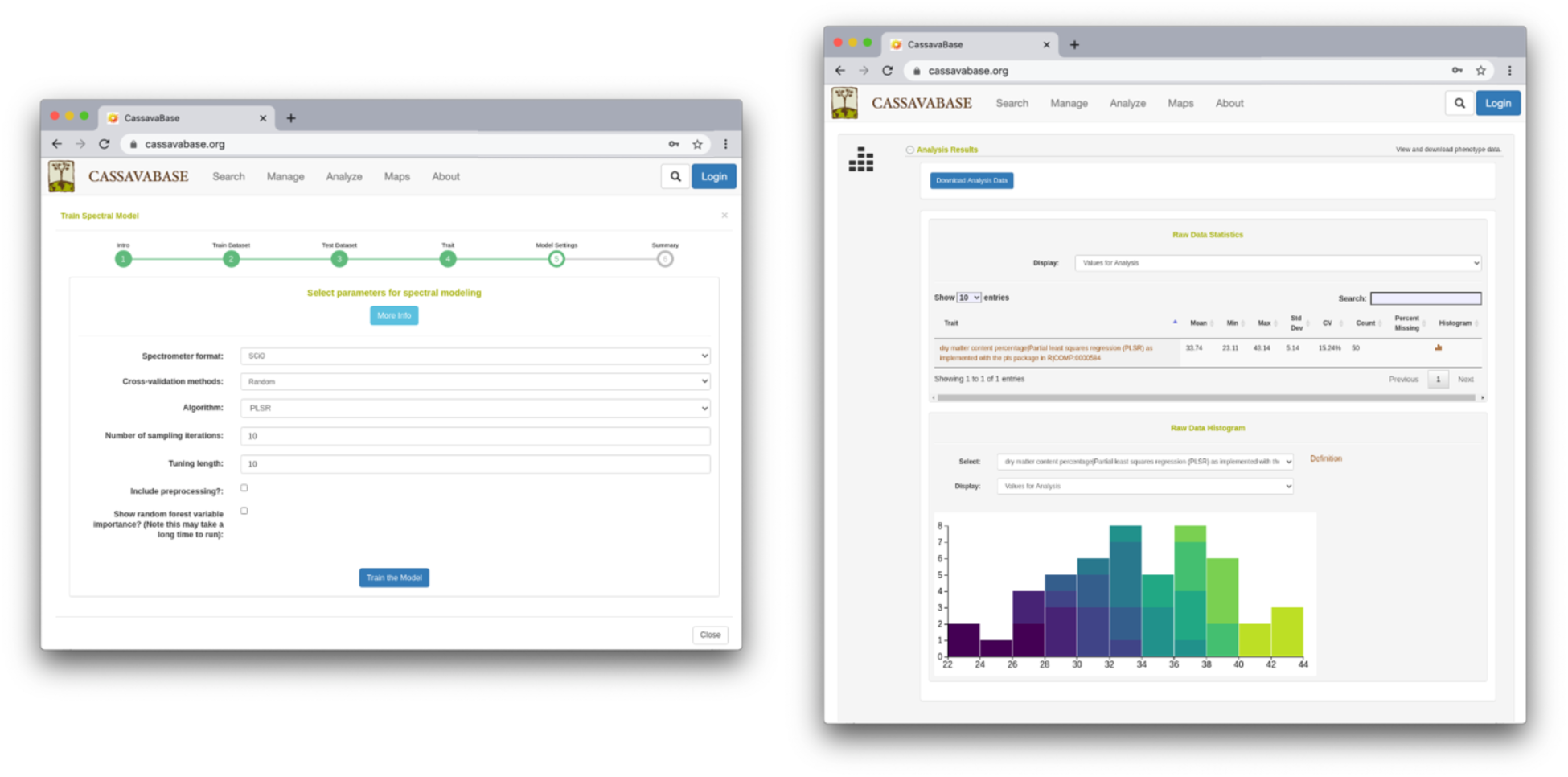
The Breedbase spectral analysis tool can be accessed using a web-based graphical user interface, shown here in the Cassavabase Breedbase instance.

**Figure 3.**
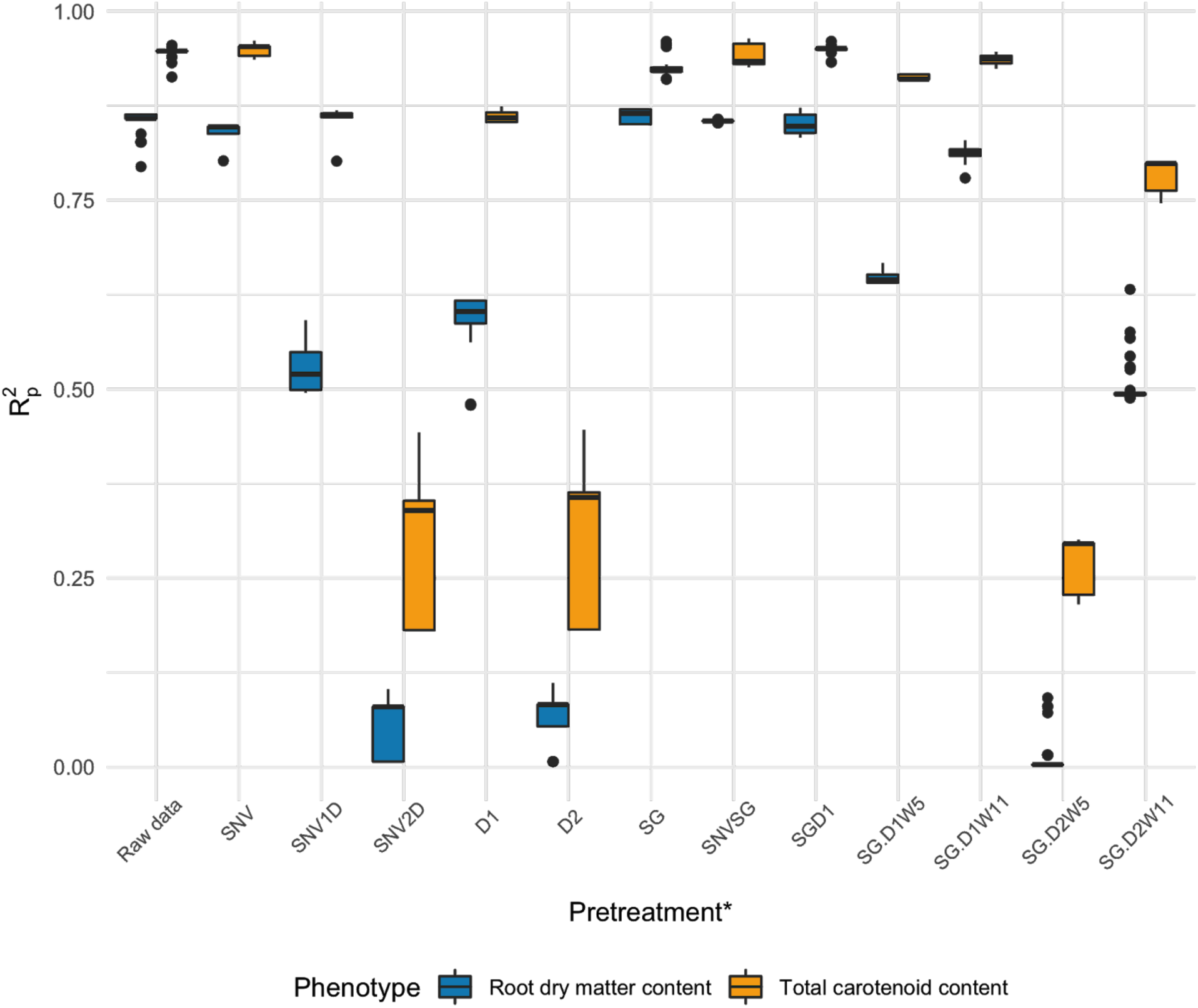
Distributions of R^2^_p_, the squared Pearson’s correlation between predicted and observed for the test set, for partial least squares regression (PLSR) models of two root quality traits trained on samples from the C16Mcal dataset and tested on samples from the C16Mval dataset from Ikeogu et al. (2017) with raw data or after pretreatment. *SNV: standard normal variate, SNV1D: standard normal variate and first derivative, SNV2D: standard normal variate and second derivative, D1: first derivative, D2: second derivative, SG: Savitzky-Golay with window size = 11, SNVSG: standard normal variate and Savitzky-Golay, SGD1: gap segment derivative with window size = 11, SG.D1W5: Savitzky-Golay with window size = 5 and first derivative, SG.D1W11: Savitzky-Golay with window size = 11 and first derivative, SG.D2W5: Savitzky-Golay with window size = 5 and second derivative, SG.D2W11: Savitzky-Golay with window size = 11 and second derivative.

## CONCLUSION

The open-source R package *waves* provides a comprehensive spectral data analysis pipeline that is tailored to the unique needs of plant breeders and their breeding programs. Integration of this package with Breedbase provides users with a GUI for improved ease-of-use within a complete digital ecosystem. Ultimately, these tools will help to improve the turnaround time for routine non-destructive phenotyping analysis, facilitating more rapid critical decision making in plant breeding programs.

## Supporting information

Supplemental Tables and User Manuals

## ACKNOWLEDGMENTS

We thank the NextGen Cassava project for their interest in and support of this work, especially Prasad Peteti and Afolabi Agbona for many fruitful discussions on ontology implementation. Thanks also to Gaby Mbanjo, Enoch Wembabazi, and Joshua Anderson for testing the pipeline. This work has been supported by USDA NIFA AFRI EWD Predoctoral Fellowship 2019-67011-29606 (J.H.), NSF BREAD IOS-1543958 (M.A.G.), the UK Foreign, Commonwealth & Development Office (L.A.M.), the Bill & Melinda Gates Foundation grant INV-007637 (L.A.M.), AfricaYam (L.A.M.), and GT4SP (L.A.M.).

## SUPPLEMENTAL MATERIAL

- Ontology table (See *Supplementary Table 1* in the Figures and Tables section)
- Table of example dataset summarized model training results (See *Supplementary Table 2* in the Figures and Tables section)
- Spectral data handling and analysis sections from Breedbase manual
- *waves* R Package documentation in CRAN format

