## Supplemental Tables and User Manuals for "Making WAVES in Breedbase: An Integrated Spectral Data Storage and Analysis Pipeline for Plant Breeding Programs"

**SUPPLEMENTAL MATERIAL**

1. Statistical ontology terms for spectroscopy analysis algorithms
2. Example dataset summarized model training results
3. Spectral data handling and analysis sections from Breedbase user manual
4. *waves* R Package documentation in CRAN format

**Supplementary Table 1. Statistical ontology terms for spectroscopy analysis algorithms**

| |
|---|
| id: SGNSTAT:0000014<br>name: Partial least squares regression (PLSR) as implemented with the pls package in R<br>is_a: SGNSTAT:0000000 |
| id: SGNSTAT:0000015<br>name: Random Forest (RF) regression as implemented with the RandomForest package in R<br>is_a: SGNSTAT:0000000 |
| id: SGNSTAT:0000016<br>name: Support vector machine (SVM) regression with linear kernel as implemented with the kernLab package in R<br>is_a: SGNSTAT:0000000 |
| id: SGNSTAT:0000017<br>name: Support vector machine (SVM) regression with radial kernel as implemented with the kernLab package in R<br>is_a: SGNSTAT:0000000 |

**Supplementary Table 2. Example dataset summarized model training results.**

| | | | **Derived from trained model only** | | **Derived from observed vs predicted values from the test set** | | | | | | | |
|---|---|---|---|---|---|---|---|---|---|---|---|---|
| **Trait** | **Pretreatment†** | **Best number of components** | **RMSECV** | **$R^2_{cv}$** | **RMSEP** | **$R^2_p$** | **RPD** | **RPIQ** | **CCC** | **Bias** | **SE** | **$R^2_{sp}$** |
| **DMC** | Raw data | 11 | 1.011 | 0.944 | 1.501 | 0.853 | 2.563 | 2.858 | 0.914 | 0.347 | 1.515 | 0.854 |
| | SNV | 11 | 0.983 | 0.947 | 1.560 | 0.842 | 2.463 | 2.746 | 0.906 | 0.370 | 1.575 | 0.850 |
| | SNV + D1 | 6 | 0.488 | 0.974 | 2.768 | 0.529 | 1.391 | 1.551 | 0.714 | 0.554 | 2.794 | 0.523 |

| | | | | | | | | | | | | |
|---|---|---|---|---|---|---|---|---|---|---|---|---|
| | SNV + D2 | 3 | 3.006 | 0.497 | 4.019 | 0.057 | 0.955 | 1.065 | 0.202 | 0.219 | 4.058 | 0.037 |
| | D1 | 6 | 0.943 | 0.950 | 2.502 | 0.599 | 1.537 | 1.714 | 0.747 | 0.633 | 2.525 | 0.598 |
| | D2 | 3 | 2.812 | 0.562 | 4.000 | 0.071 | 0.960 | 1.071 | 0.238 | 0.264 | 4.038 | 0.048 |
| | SG (W11) | 13 | 1.030 | 0.942 | 1.469 | 0.860 | 2.616 | 2.916 | 0.918 | 0.353 | 1.483 | 0.872 |
| | SNV + SG | 11 | 1.012 | 0.944 | 1.499 | 0.855 | 2.561 | 2.855 | 0.915 | 0.371 | 1.514 | 0.861 |
| | GD | 10 | 0.980 | 0.947 | 1.522 | 0.849 | 2.525 | 2.816 | 0.911 | 0.338 | 1.537 | 0.861 |
| | SG + D1 (W5) | 16 | 0.294 | 0.991 | 2.328 | 0.646 | 1.650 | 1.839 | 0.796 | 0.371 | 2.350 | 0.671 |
| | SG + D1 (W11) | 9 | 0.725 | 0.970 | 1.682 | 0.814 | 2.284 | 2.547 | 0.891 | 0.368 | 1.698 | 0.810 |
| | SG + D2 (W5) | 1 | 3.286 | 0.371 | 4.141 | 0.011 | 0.928 | 1.034 | 0.069 | 0.247 | 4.180 | 0.011 |
| | SG + D2 (W11) | 25 | 0.201 | 0.990 | 2.811 | 0.504 | 1.367 | 1.525 | 0.703 | 0.331 | 2.838 | 0.490 |
| | Raw data | 13 | 1.191 | 0.976 | 1.743 | 0.946 | 4.306 | 7.644 | 0.973 | 0.114 | 1.760 | 0.897 |
| | SNV | 10 | 1.218 | 0.976 | 1.662 | 0.950 | 4.532 | 8.046 | 0.974 | 0.159 | 1.678 | 0.903 |
| | SNV + D1 | 8 | 1.077 | 0.979 | 2.785 | 0.862 | 2.692 | 4.778 | 0.927 | 0.223 | 2.811 | 0.738 |
| | SNV + D2 | 1 | 5.458 | 0.501 | 6.257 | 0.304 | 1.202 | 2.135 | 0.484 | 0.437 | 6.316 | 0.266 |
| | D1 | 6 | 1.148 | 0.977 | 2.804 | 0.860 | 2.672 | 4.744 | 0.926 | 0.218 | 2.830 | 0.743 |
| | D2 | 3 | 5.329 | 0.525 | 6.215 | 0.315 | 1.210 | 2.147 | 0.497 | 0.566 | 6.273 | 0.272 |
| **TCC** | SG (W11) | 29 | 0.461 | 0.993 | 2.081 | 0.926 | 3.654 | 6.487 | 0.961 | 0.212 | 2.101 | 0.867 |
| | SNV + SG | 8 | 0.869 | 0.984 | 1.789 | 0.942 | 4.272 | 7.584 | 0.970 | 0.125 | 1.805 | 0.893 |
| | GD | 10 | 1.386 | 0.968 | 1.673 | 0.950 | 4.484 | 7.960 | 0.974 | 0.093 | 1.689 | 0.899 |
| | SG + D1 (W5) | 8 | 1.007 | 0.983 | 2.194 | 0.913 | 3.415 | 6.063 | 0.955 | 0.020 | 2.214 | 0.846 |
| | SG + D1 (W11) | 11 | 1.028 | 0.982 | 1.899 | 0.936 | 3.955 | 7.022 | 0.967 | -0.070 | 1.917 | 0.874 |
| | SG + D2 (W5) | 1 | 5.808 | 0.436 | 6.479 | 0.270 | 1.158 | 2.055 | 0.439 | -0.535 | 6.539 | 0.257 |
| | SG + D2 (W11) | 9 | 1.228 | 0.975 | 3.460 | 0.785 | 2.167 | 3.848 | 0.880 | 0.332 | 3.493 | 0.706 |

Summary statistics for 50 iterations of PLSR for root dry matter content (DMC) and total carotenoid content (TCC) prediction with 5-fold cross-validation for hyperparameter tuning. The C16Mcal (*n* = 120) dataset was used for training and the C16Mval (*n* = 53) dataset was used for testing, both from Ikeogu et al. (2017).

†SNV: standard normal variate, SNV + 1D: standard normal variate and first derivative, SNV + 2D: standard normal variate and second derivative, D1: first derivative, D2: second derivative, SG: Savitzky-Golay with window size = 11, SNV + SG: standard normal variate and Savitzky-Golay, GD: gap segment derivative with window size = 11, SG + D1 (W5): Savitzky-Golay with window size = 5 and first derivative, SG + D1 (W11): Savitzky-Golay with window size = 11 and first derivative, SG + D2 (W5): Savitzky-Golay with window size = 5 and second derivative, SG + D2 (W11): Savitzky-Golay with window size = 11 and second derivative.

Reported statistics:

- Best number of components, the best number of components to be included in a PLSR model
- RMSECV, the root mean squared error of cross-validation
- $R^2_{cv}$, the coefficient of multiple determination of cross-validation for PLSR models
- RMSEP, the root mean squared error of prediction
- $R^2_{p}$, the squared Pearson's correlation between predicted and observed test set values
- RPD, the ratio of standard deviation of observed test set values to RMSEP
- RPIQ, the ratio of performance to interquartile difference
- CCC, the concordance correlation coefficient
- Bias, the average difference between the predicted and observed values
- SE, the standard error of prediction
- $R^2_{sp}$, the squared Spearman's rank correlation between predicted and observed test set values

**Supplemental file 1**: Breedbase spectral tool manual

#### Breedbase spectral data upload and storage

Visible and near-infrared spectroscopy (vis-NIRS) can be related to reference phenotypes through statistical models to produce accurate phenotypic predictions for unobserved samples, increasing phenotyping throughput. This technique is commonly used for predicting traits such as total starch, protein, carotenoid, and water content in many plant breeding programs. Breedbase implements the R package *waves* to offer training, evaluation, storage, and use of vis-NIRS prediction models for a wide range of spectrometers and phenotypes.

Spectral data handling in Breedbase makes use of the R package *waves* for tasks such as outlier identification, plotting, sample aggregation, model calibration, and trait prediction.

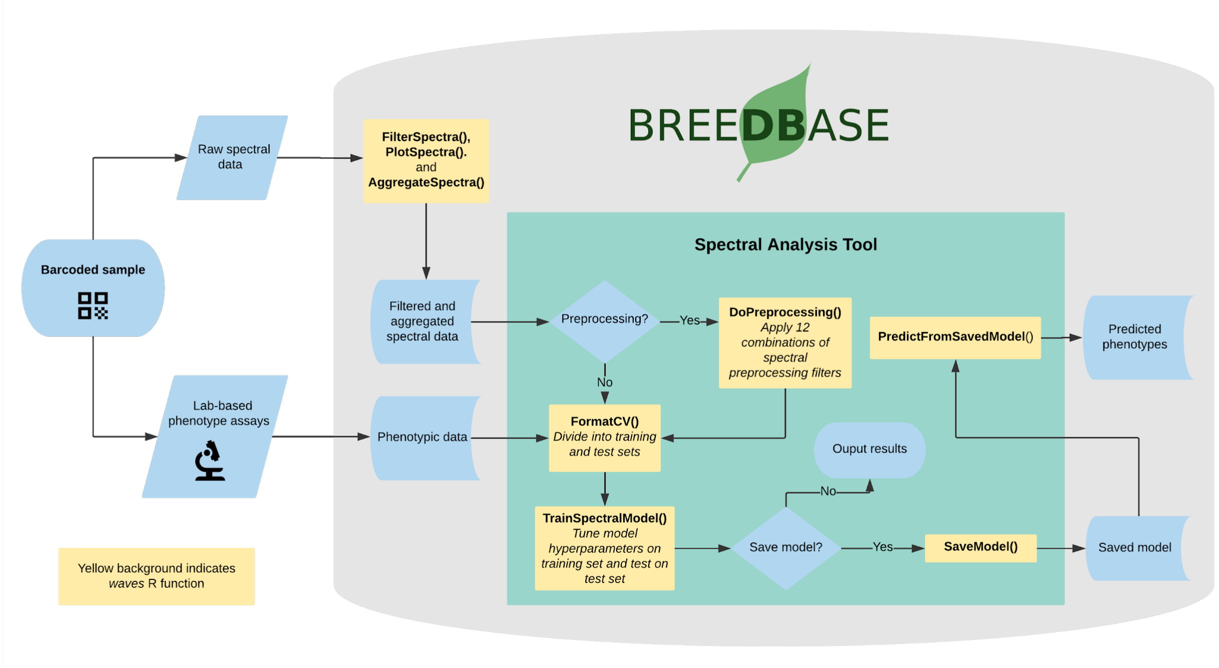

#### Ontology

Breedbase instances are governed by controlled ontologies based on the Crop Ontology system (Shrestha et al., 2012). We have extended the statistical ontology for this system to include spectroscopy analysis algorithm terms, and these terms are combined with existing trait ontology terms to generate spectroscopy-based predicted ontology terms.

| |
|---|
| id: SGNSTAT:0000014<br>name: Partial least squares regression (PLSR) as implemented with the pls package in R<br>is_a: SGNSTAT:0000000 |
| id: SGNSTAT:0000015<br>name: Random Forest (RF) regression as implemented with the RandomForest package in R<br>is_a: SGNSTAT:0000000 |

| |
|---|
| id: SGNSTAT:0000016<br>name: Support vector machine (SVM) regression with linear kernel as implemented with the kernLab package in R<br>is_a: SGNSTAT:0000000 |
| id: SGNSTAT:0000017<br>name: Support vector machine (SVM) regression with radial kernel as implemented with the kernLab package in R<br>is_a: SGNSTAT:0000000 |

**Upload**
Breedbase has implemented a flexible spectral data storage protocol that handles spectral data irrespective of the source spectrometer. Spectral data can be uploaded as a CSV file that includes metadata in the leftmost columns followed by one column per spectral measurement to the right. Rows represent a single scan or sample, each with a unique ID that must match to a Breedbase observationUnitName. Future data transfer using BrAPI will allow for interoperability with data collection software.

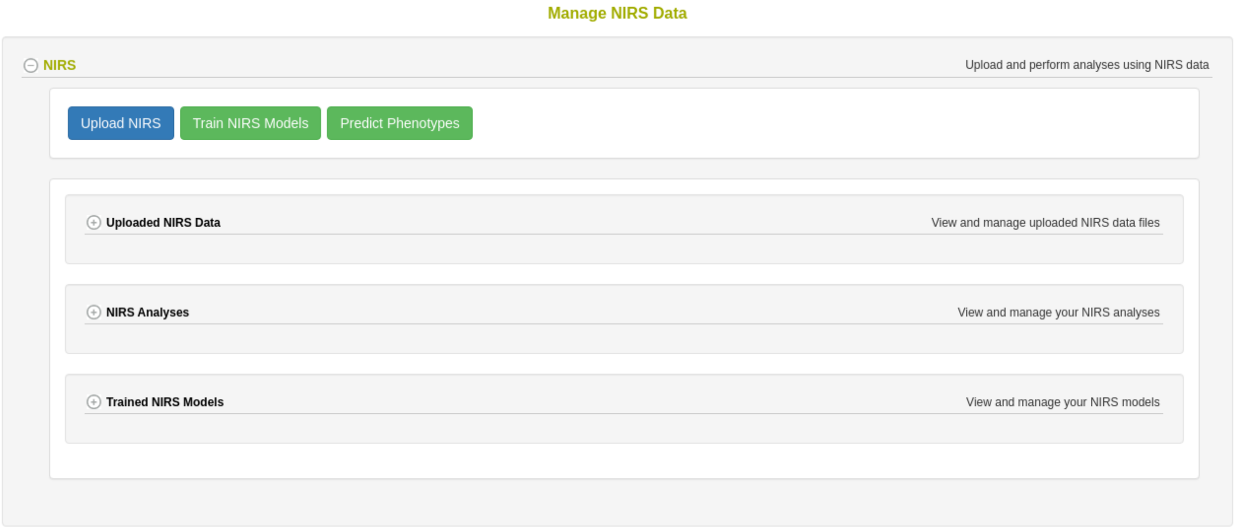

| id | sample_id | sampling_date | observationunit_name | device_id | device_type | comments | 740 | 741 | 742 | 743 | 744 | 745 | 746 |
|---|---|---|---|---|---|---|---|---|---|---|---|---|---|
| 1 | 7a6ac477-d291-4d07-a6 | 2020-6-24 | myTrial20_rep1_acc_001 | 503E4BFC4E923999 | SCIO | | 0.885707958 | 0.88572938 | 0.885590265 | 0.885493457 | 0.885493162 | 0.885572662 | 0.885628732 |
| 2 | 7a6ac477-d291-4d07-a6 | 2020-6-24 | myTrial20_rep1_acc_002 | 503E4BFC4E923999 | SCIO | | 0.909132994 | 0.908724223 | 0.90824451 | 0.907891706 | 0.907697281 | 0.907626115 | 0.90755938 |
| 3 | 7a6ac477-d291-4d07-a6 | 2020-6-24 | myTrial20_rep1_acc_003 | 503E4BFC4E923999 | SCIO | | 0.889220207 | 0.889013119 | 0.888681812 | 0.888431257 | 0.888310362 | 0.888297321 | 0.888284052 |
| 4 | 73c648ca-f5fb-4231-a18 | 2020-6-24 | myTrial20_rep1_acc_004 | 503E4BFC4E923999 | SCIO | | 0.8900087 | 0.889604969 | 0.889191073 | 0.888958654 | 0.88893379 | 0.889072741 | 0.8892472 |
| 5 | 73c648ca-f5fb-4231-a18 | 2020-6-24 | myTrial20_rep1_acc_005 | 503E4BFC4E923999 | SCIO | | 0.939101707 | 0.93868202 | 0.93820132 | 0.937873867 | 0.937742819 | 0.937775129 | 0.93784594 |
| 6 | 73c648ca-f5fb-4231-a18 | 2020-6-24 | myTrial20_rep1_acc_006 | 503E4BFC4E923999 | SCIO | | 0.876289461 | 0.875981159 | 0.875579263 | 0.875289805 | 0.875162225 | 0.875171404 | 0.87520442 |
| 7 | d5b55c93-4c8e-4ef6-9c | 2020-6-24 | myTrial20_rep1_acc_007 | 503E4BFC4E923999 | SCIO | | 0.879217838 | 0.878925781 | 0.878588441 | 0.8783872 | 0.878346547 | 0.878423929 | 0.878495739 |
| 8 | 7a6ac477-d291-4d07-a6 | 2020-6-24 | myTrial20_rep1_acc_008 | 503E4BFC4E923999 | SCIO | | 0.890746588 | 0.890515672 | 0.89016542 | 0.889903562 | 0.889783304 | 0.889782828 | 0.889792154 |
| 9 | 7a6ac477-d291-4d07-a6 | 2020-6-24 | myTrial20_rep1_acc_009 | 503E4BFC4E923999 | SCIO | | 0.850444238 | 0.85032422 | 0.850094039 | 0.8499495 | 0.849942163 | 0.850052119 | 0.850175036 |

**Evaluate and remove spectral outliers**
Spectral calibration models can be heavily affected by the presence of outliers, whether they come from spectrometer spectral artifacts or user errors. Mahalanobis distance (Mahalanobis, 1936) is a measure of the distance between a single observation and a larger distribution and is commonly used in the identification of outliers in a multivariate space (De Maesschalck et al., 2000). The *FilterSpectra()* function in *waves* calculates the Mahalanobis distance of each

observation in a given spectral matrix using the *mahalanobis()* function from the *stats* package. Observations are identified as outliers if the squared distance is greater than the 95th percentile of a $X^2$-distribution with p degrees of freedom, where p is the number of columns (wavelengths) in the spectral matrix (Johnson and Wichern, 2007). In Breedbase, this procedure is applied on a per-dataset basis on upload and outliers are given binary tags "Outlier."

**Plot spectra with and without outliers highlighted**

After outlier identification, a plot is generated using the PlotSpectra() function in *waves*. This function uses the filtered spectra and the *ggplot()* function from the *ggplot2* package to create a line plot with outliers highlighted by color. A list of rows identified as outliers are shown beneath the plot. Plots are saved as .png files and linked to the original input datasets. Plot image files can be downloaded via the "Download Plot" button on the upload webpage.

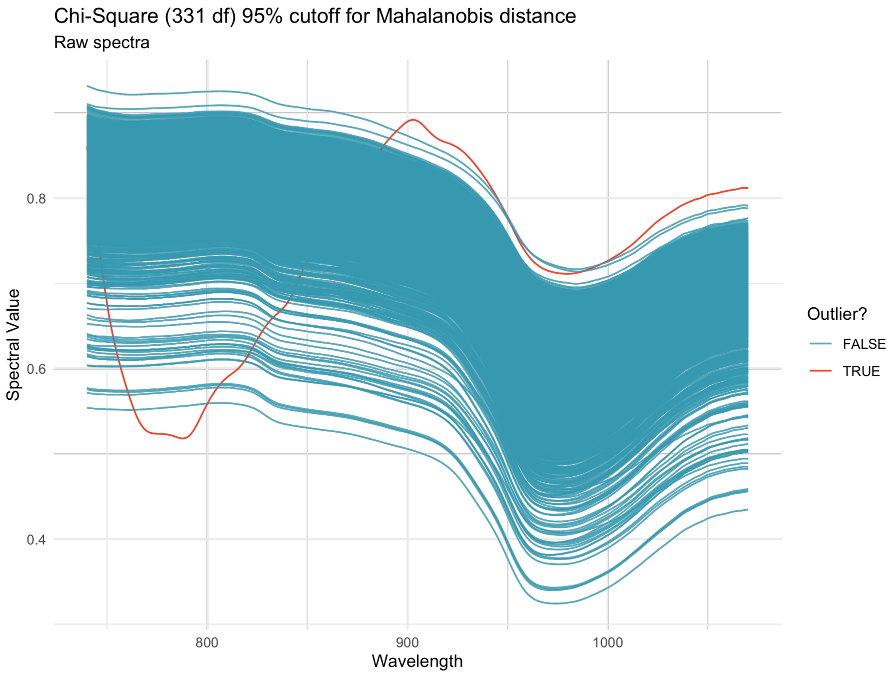

Outlier identification

**Aggregate spectra by observation unit**

To obtain a stable and reliable spectral profile, most spectrometer manufacturers recommend that multiple spectral scans are captured for each sample. While some spectrometers aggregate these scans internally, many do not, requiring the user to do so before analysis can take place. Breedbase handles these cases upon data upload following filtering steps by calling the *AggregateSpectra()* function from *waves*, saving the aggregated scans for future access through the search wizard feature. Scans are aggregated by sample mean (e.g., plot-level basis) according to the provided observationUnitName field. The aggregated, representative scan for each observationUnit is accessible through the search wizard feature on Breedbase. After aggregation, the user exits the upload workflow and the raw data file is saved in the upload archive.

**Breedbase Spectral Analysis Tool Manual**

Visible and near-infrared spectroscopy (vis-NIRS) can be related to reference phenotypes through statistical models to produce accurate phenotype predictions for unobserved samples, increasing phenotyping throughput. This technique is commonly used for predicting traits such as total starch, protein, carotenoid, and water content in many plant breeding programs. Breedbase implements the R package *waves* to offer training, evaluation, storage, and use of vis-NIRS prediction models for a wide range of spectrometers and phenotypes.

**Dataset selection**

In order to initiate an analysis, the user must select one or more datasets using the Breedbase search wizard tool. A dataset in Breedbase can contain observationUnit-level (plot-, plant-, or sample-level) trial metadata and phenotypic data from one or more trials. After navigating to the "Spectral Analysis" webpage under the "Analysis" tab in Breedbase, the user can enter the analysis workflow and select one of these datasets as input for model training. An optional test dataset can be selected in the second step of the workflow.

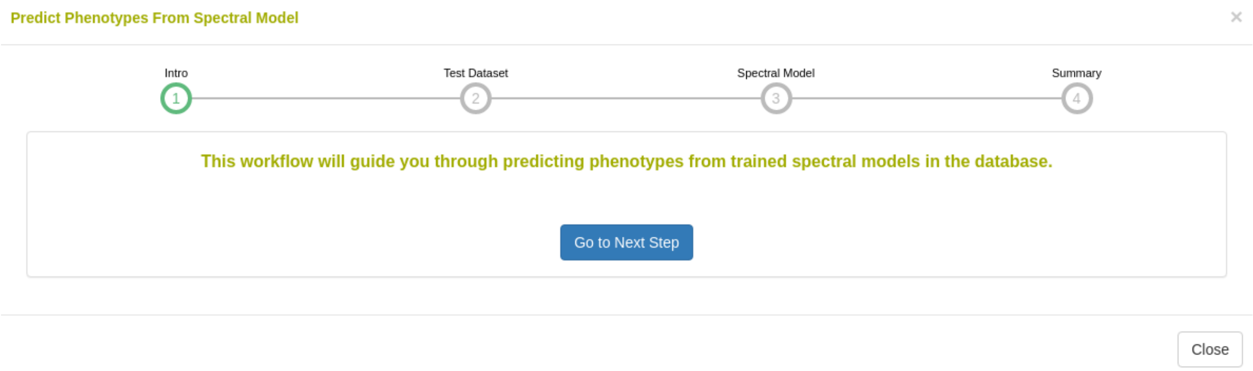

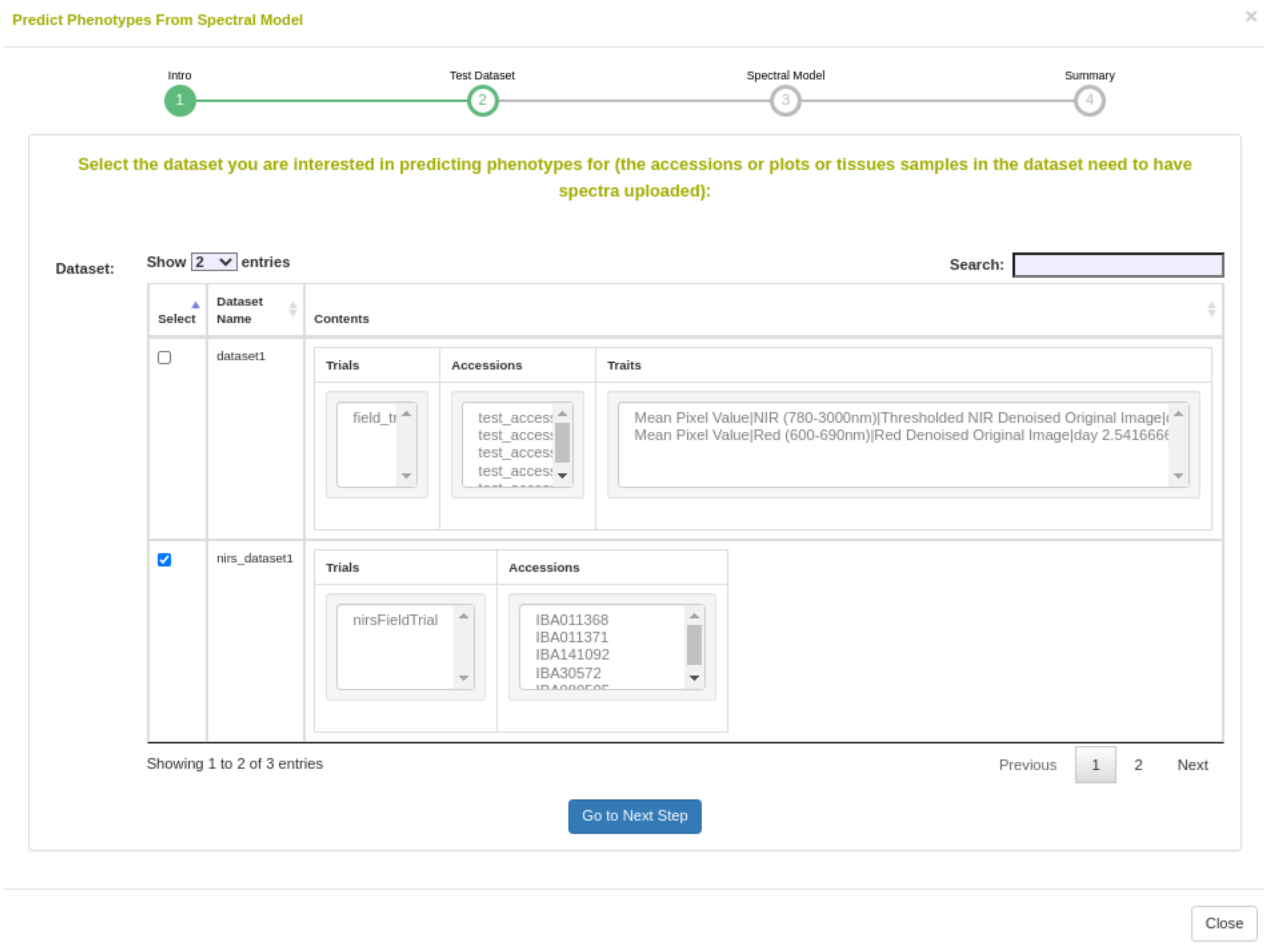

#### Cross-validation

Five cross-validation schemes that represent scenarios common in plant breeding are available for this analysis. These include CV1, CV2, CV0, and CV00 as outlined below and described in depth by Jarquín et al. (2017) as well as random and stratified random sampling. For those schemes from Jarquín et al. (2017), specific input datasets must be chosen based on genotype and environment relatedness.

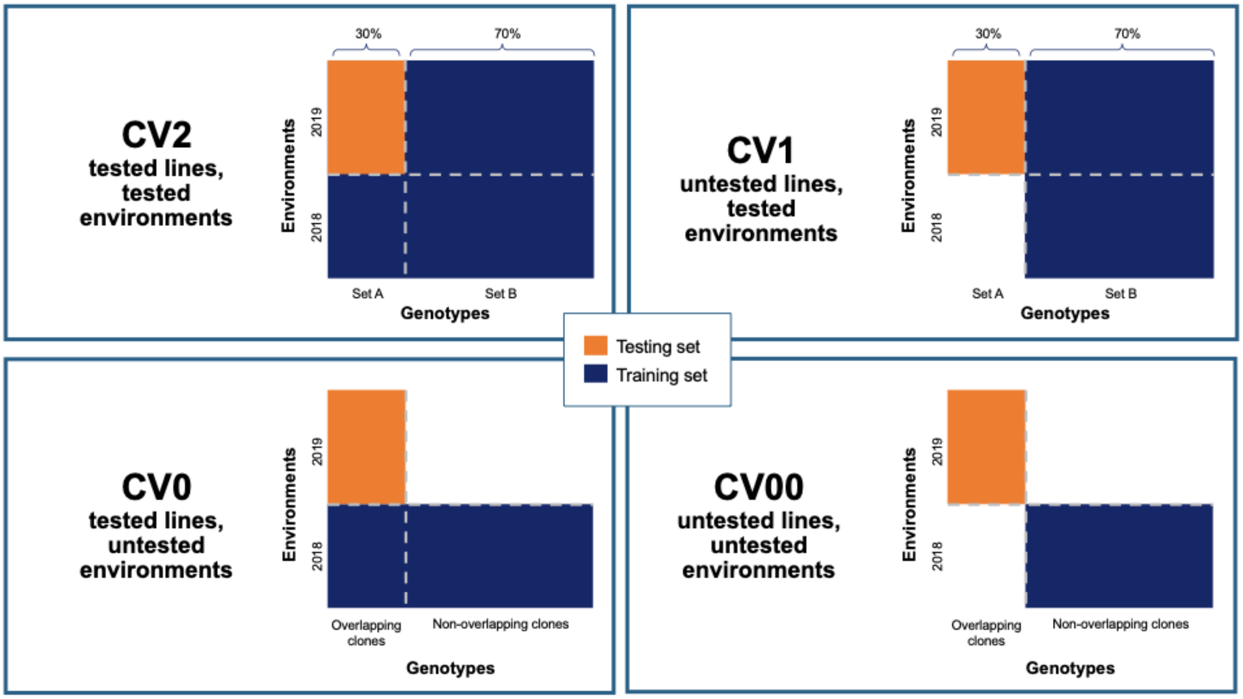

**Preprocessing**

Preprocessing, also known as pretreatment, is often used to increase the signal to noise ratio in vis-NIR datasets. The *waves* function *DoPreprocessing()* applies functions from the *stats* and *prospectr* packages for common spectral preprocessing methods with the following options:

- Raw data (default)
- First derivative
- Second derivative
- Gap segment derivative
- Standard normal variate (SNV; Barnes et al., 1989)
- Savitzky-Golay polynomial smoothing (Savitzky and Golay, 1964)

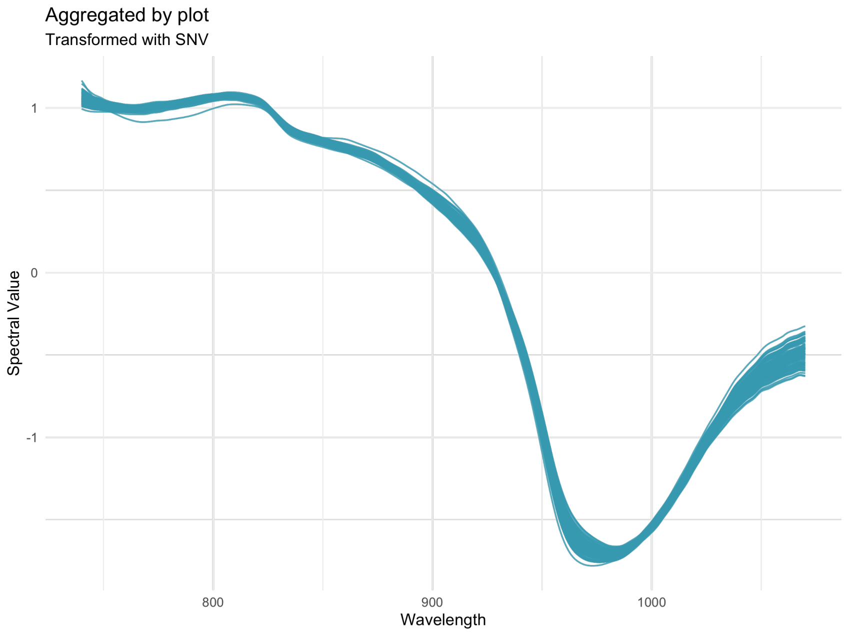

Spectra pretreated with the SNV filter

**Algorithms**

Several algorithms are available for calibration model development in Breedbase via the *waves* package. The *TrainSpectralModel()* function in *waves* performs hyperparameter tuning as applicable using these algorithms in combination with cross validation and train functions from the package *caret*. Currently, only regression algorithms are available, but classification algorithms such as PLS-DA and SVM classification are under development.

- Partial least squares regression (PLSR; Wold et al., 1982; Wold et al., 1984) is a popular method for spectral calibrations, as it can handle datasets with high levels of collinearity, reducing the dimensionality of these data into orthogonal latent variables ('components') that are then related to the response variable through a linear model (reviewed in Wold et al., 2001). To avoid overfitting, the number of these components included in the final model must be tuned for each use case. The PLSR algorithm from the *pls* package is implemented by *waves*.
- Random Forest regression (RF; Ho, 1995) is a machine learning algorithm based on a series of decision trees. The number of trees and decisions at each junction are hyperparameters that must be tuned for each model. Another feature of this algorithm is the ability to extract variable importance measures from a fitted model (Breiman, 2001). In Breedbase, this option is made available through implementation of the RF algorithm from the package *randomForest* in the *waves* function *TrainSpectralModel()*. This function outputs both model performance statistics and a downloadable table of importance values for each wavelength. It is worth noting that this algorithm is computationally intensive, so the user should not be alarmed if results do not come right away. Breedbase will continue to work in the background and will display results when the analysis is finished.
- Support vector machine regression (SVM; Vapnik, 2000) is another useful algorithm for working with high-dimensional datasets consisting of non-linear data, with applications

in both classification and regression. The package *waves* implements SVM with both linear and radial basis function kernels using the *kernlab* package.

**Output common model summary statistics**

After training, model performance statistics are both displayed on a results webpage and made available for download in .csv format. These statistics are calculated by the *TrainSpectralModel()* function in *waves* using the *caret* and *spectacles* packages. Reported statistics include:

- Tuned parameters depending on the model algorithm
    - Best.n.comp, the best number of components to be included in a PLSR model
    - Best.ntree, the best number of trees in an RF model
    - Best.mtry, the best number of variables to include at every decision point in an RF model
- RMSECV, the root mean squared error of cross-validation
- $R^2_{cv}$, the coefficient of multiple determination of cross-validation for PLSR models
- RMSEP, the root mean squared error of prediction
- $R^2_p$, the squared Pearson's correlation between predicted and observed test set values
- RPD, the ratio of standard deviation of observed test set values to RMSEP
- RPIQ, the ratio of performance to interquartile difference
- CCC, the concordance correlation coefficient
- Bias, the average difference between the predicted and observed values
- SEP, the standard error of prediction
- $R^2_{sp}$, the squared Spearman's rank correlation between predicted and observed test set values

**Export model for later use**

Once a model has been trained, it can be stored for later use. This action calls the *SaveModel()* function from *waves*. Metadata regarding the training dataset and other parameters specified by the user upon training initialization are stored alongside the model object itself in the database.

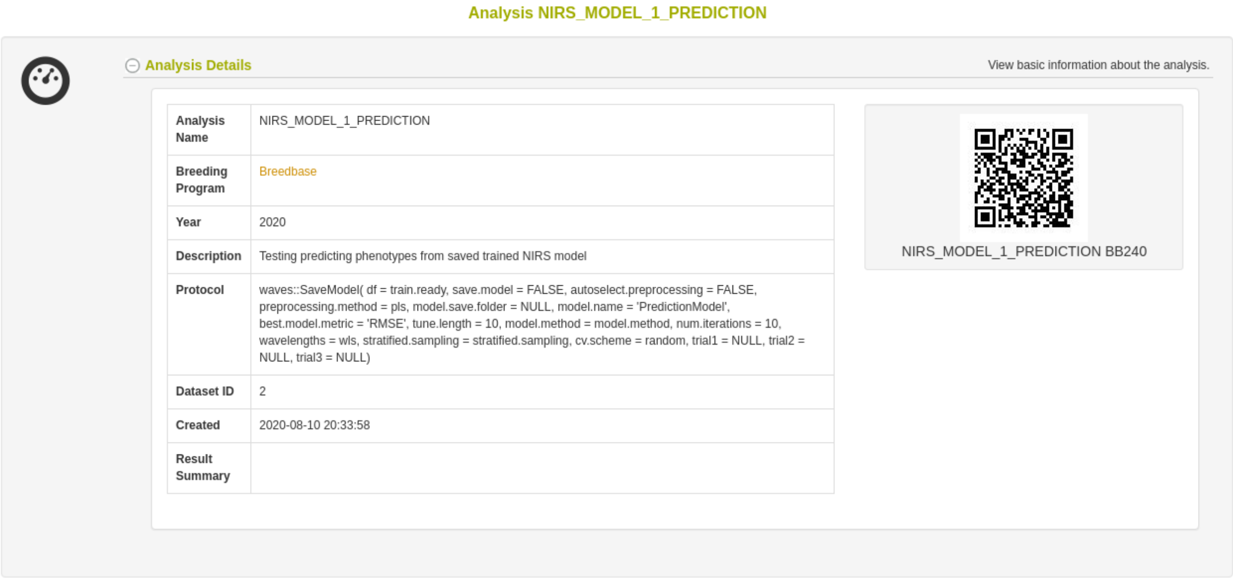
Analysis NIRS_MODEL_1_PREDICTION

Analysis Details

View basic information about the analysis.

| | |
|---|---|
| Analysis Name | NIRS_MODEL_1_PREDICTION |
| Breeding Program | Breedbase |
| Year | 2020 |
| Description | Testing predicting phenotypes from saved trained NIRS model |
| Protocol | waves::SaveModel( df = train.ready, save.model = FALSE, autoselect.preprocessing = FALSE, preprocessing.method = pls, model.save.folder = NULL, model.name = 'PredictionModel', best.model.metric = 'RMSE', tune.length = 10, model.method = model.method, num.iterations = 10, wavelengths = wls, stratified.sampling = stratified.sampling, cv.scheme = random, trial1 = NULL, trial2 = NULL, trial3 = NULL) |
| Dataset ID | 2 |
| Created | 2020-08-10 20:33:58 |
| Result Summary | |

NIRS_MODEL_1_PREDICTION BB240

**Predict phenotypes from an exported model (routine use)**

For phenotype predictions, users can select the model of their choice, and if their selected dataset has scans that were taken using the same spectrometer model, the phenotype the model was trained on can be used for prediction. Predicted phenotypes are stored as such in the database and are tagged with an ontology term specifying that they are predicted and not directly measured. Metadata regarding the model used for prediction is stored alongside the predicted value in the database. Predicted phenotypes can then be used as normal in other Breedbase analysis tools such as the Selection Index and GWAS.

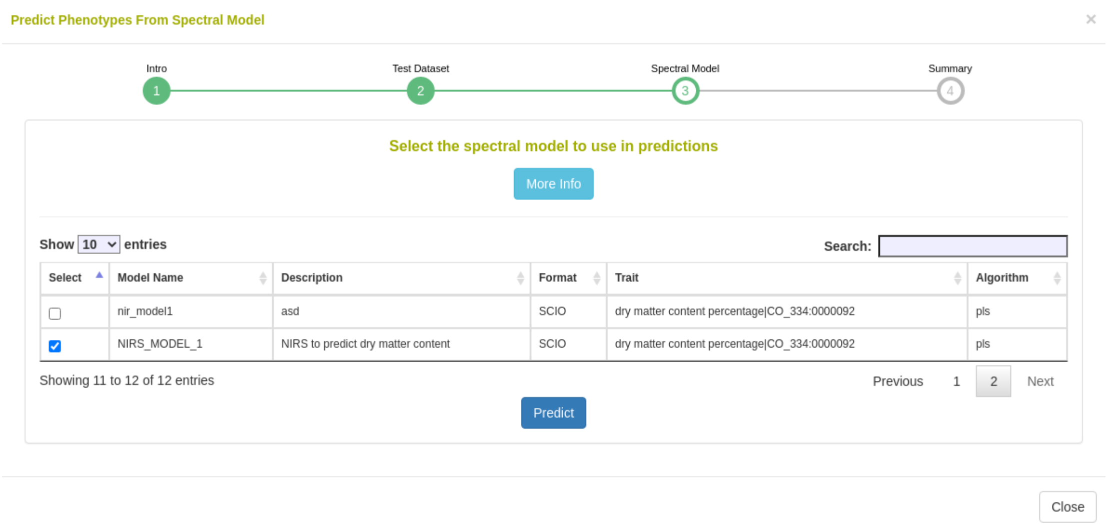

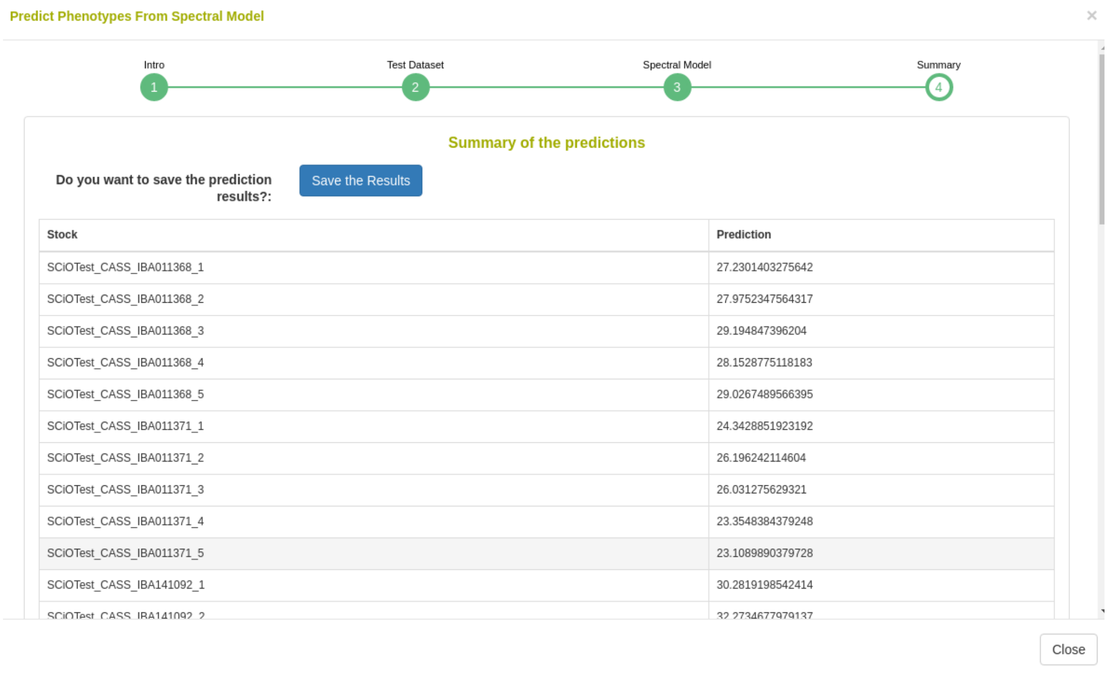

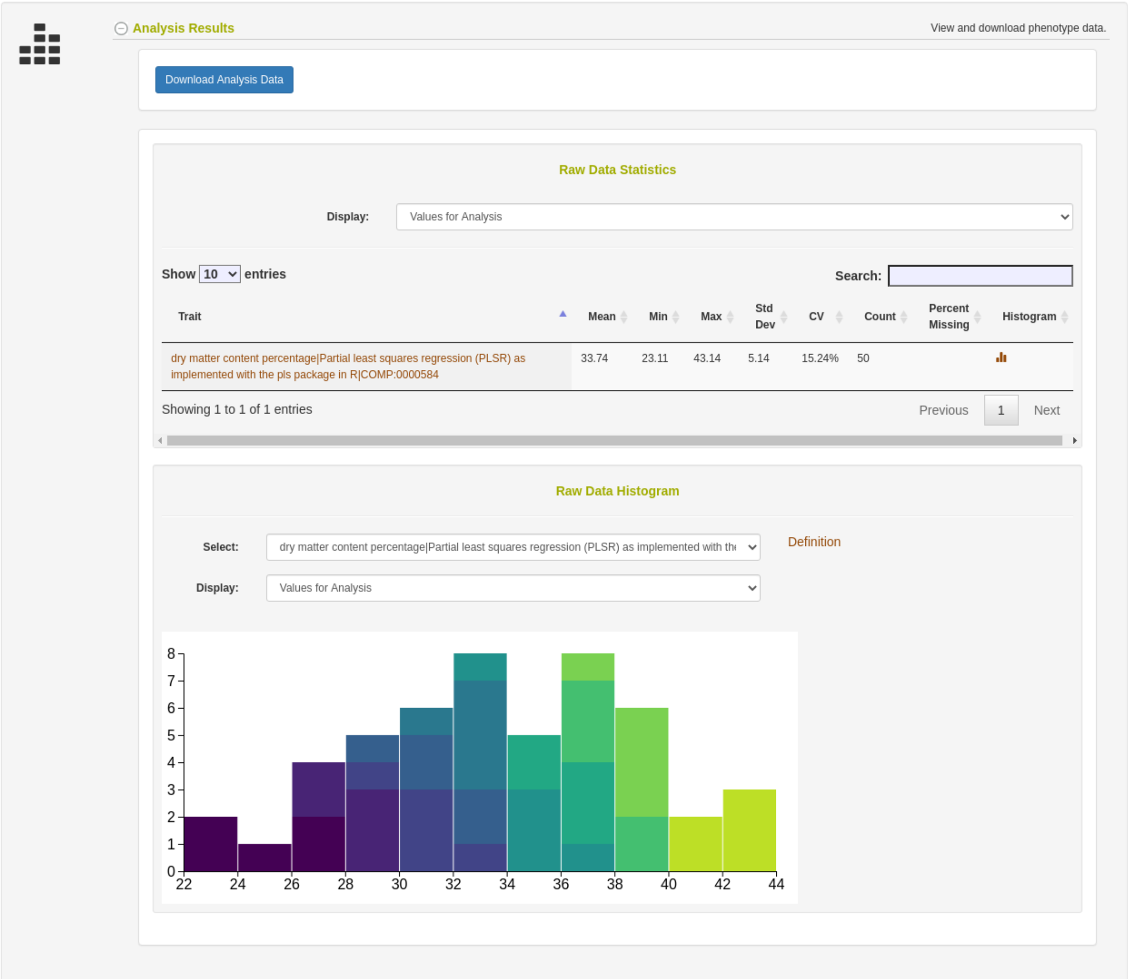

#### FAQ Storage

- Though Breedbase does not handle proprietary data formats, once they are transformed into CSV files, the number of wavelengths and size of gaps between them does not limit compatibility with Breedbase.

#### FAQ Analysis

- Breedbase does not allow prediction models involving data from multiple spectrometers types at once.

**Supplemental File 2. waves R package documentation**

### Package ‘waves’

September 17, 2020

**Title** Vis-NIR Spectral Analysis Wrapper

**Version** 0.1.0

**Description** Originally designed application in the context of resource-limited plant research and breeding programs, 'waves' provides an open-source solution to spectral data processing and model development by bringing useful packages together into a streamlined pipeline. This package is wrapper for functions related to the analysis of point visible and near-infrared reflectance measurements. It includes visualization, filtering, aggregation, preprocessing, cross-validation set formation, model training, and prediction functions to enable open-source association of spectral and reference data.
Specialized cross-validation schemes are described in detail in Jarquín et al. (2017) <doi:10.3835/plantgenome2016.12.0130>. Example data is from Ikeogu et al. (2017) <doi:10.1371/journal.pone.0188918>.

**URL** https://github.com/GoreLab/waves

**Maintainer** Jenna Hershberger <>

**BugReports** https://github.com/GoreLab/waves/issues

**Depends** R (>= 3.5)

**License** MIT + file LICENSE

**Encoding** UTF-8

**LazyData** true

**Imports** dplyr, prospectr, spectacles, caret, pls, randomForest, wesanderson, magrittr, tidyselect, ggplot2, tidyr (>= 1.0), stringr, rlang

**RoxygenNote** 6.1.1

**Suggests** testthat (>= 2.1.0)

**NeedsCompilation** no

**Author** Jenna Hershberger [aut, cre] (<https://orcid.org/0000-0002-3147-6867>), Michael Gore [ths], NSF BREAD IOS-1543958 [fnd]

**Repository** CRAN

**Date/Publication** 2020-09-17 12:10:02 UTC

### R topics documented:

**Index** **17**

---

`AggregateSpectra` *Aggregate data based on grouping variables and a user-provided function*

---

##### Description

Use grouping variables to collapse spectral `data.frame` by mean or median. Recommended for use after `FilterSpectra`

##### Usage

```
AggregateSpectra(df, grouping.colnames, reference.value.colname,
  agg.function)
```

##### Arguments

`df` — `data.frame` object containing one or multiple columns of grouping variables (must be consistent within each group), column of reference values (optional), and columns of spectra. Spectral column names must start with "X".

`grouping.colnames` — Names of columns to be used as grouping variables. Minimum 2 variables required. Default is c("trial", "plot").

`reference.value.colname` — Name of reference column to be aggregated along with spectra. Default is "reference"

`agg.function` — Name of function (string format) to be used for sample aggregation. Must be either "mean" or "median". Default is "mean".

##### Value

`data.frame` object `df` aggregated based on grouping column by `agg.function`

##### Author(s)

Jenna Hershberger <>

#### Examples

```
library(magrittr)
aggregated.test <- ikeogu.2017 %>%
  dplyr::select(-TCC) %>%
  na.omit() %>%
  AggregateSpectra(grouping.colnames = c("study.name"),
                   reference.value.colname = "DMC.oven",
                   agg.function = "mean")
aggregated.test[1:5, 1:5]
```

---

`DoPreprocessing` *Preprocess spectral data according to user-designated method*

---

#### Description

Preprocessing, also known as pretreatment, is often used to increase the signal to noise ratio in vis-NIR datasets. The *waves* function `DoPreprocessing` applies common spectral preprocessing methods such as standard normal variate and the Savitzky-Golay filter.

#### Usage

```
DoPreprocessing(df, test.data = NULL, preprocessing.method = 1,
  wavelengths = 740:1070)
```

#### Arguments

`df` `data.frame` object containing spectral data. First column(s) (optional) include metadata (with or without reference value column) followed by spectral columns. Spectral column names must be formatted as "X" followed by wavelength Include no other columns to right of spectra! No missing values permitted.

`test.data` `data.frame` object with same format as train.data. Will be appended to `df` during preprocessing so that the same transformations are applied to each row. Default is `NULL`.

`preprocessing.method` Number or list of numbers 1:13 corresponding to desired pretreatment method(s):

- 1 = raw data (default)
- 2 = standard normal variate (SNV)
- 3 = SNV and first derivative
- 4 = SNV and second derivative
- 5 = first derivative
- 6 = second derivative
- 7 = Savitzky–Golay filter (SG)
- 8 = SNV and SG
- 9 = gap segment derivative (window size = 11)
- 10 = SG and first derivative (window size = 5)

- 11 = SG and first derivative (window size = 11)
- 12 = SG and second derivative (window size = 5)
- 13 = SG and second derivative (window size = 11)

`wavelengths` List of wavelengths represented by each column in `df`. Default is 740:1070.

#### Value

Preprocessed `df`‘ (or list of `data.frames`) with reference column intact

#### Author(s)

Jenna Hershberger <>

#### Examples

```
DoPreprocessing(df = ikeogu.2017, wavelengths = 350:2500)[1:5,1:5]
```

---

`FilterSpectra` *Filter spectral data frame based on Mahalanobis distance*

---

#### Description

Determine Mahalanobis distances of observations (rows) within a given `data.frame` with spectral data. Option to filter out observations based on these distances.

#### Usage

```
FilterSpectra(df, filter, return.distances, num.col.before.spectra,
  window.size, verbose)
```

#### Arguments

`df` a `data.frame` object containing columns of spectra and rows of observations. May also contain columns of metadata to the left of the spectra.

`filter` boolean that determines whether or not the input `data.frame` will be filtered. If `TRUE`, `df` will be filtered according to squared Mahalanobis distance with a 95% cutoff from a chi-square distribution with degrees of freedom = number of spectral columns. If `FALSE`, a column of squared Mahalanobis distances `h.distance` will be added to the right side of df and all rows will be returned. Default is `TRUE`.

`return.distances` boolean that determines whether a column of squared Mahalanobis distances will be included in output `data.frame`. If `TRUE`, a column of Mahalanobis distances for each row will be added to the right side of `df`. Default is `FALSE`.

`num.col.before.spectra` number of columns to the left of the spectral matrix in `df`. Default is 4.

`window.size` number defining the size of window to use when calculating the covariance of the spectra (required to calculate Mahalanobis distance). Default is 10.

verbose — If TRUE, the number of rows removed through filtering will be printed to the console. Default is TRUE.

#### Details

This function uses a chi-square distribution with 95% cutoff where degrees of freedom = number of wavelengths (columns) in the input data.frame.

#### Value

If filter is TRUE, returns filtered data frame df and reports the number of rows removed. The Mahalanobis distance with a cutoff of 95% of chi-square distribution (degrees of freedom = number of wavelengths) is used as filtering criteria. If filter is FALSE, returns full input df with column h.distances containing the Mahalanobis distance for each row.

#### Author(s)

Jenna Hershberger <>

#### Examples

```
library(magrittr)
filtered.test <- ikeogu.2017 %>%
  dplyr::select(-TCC) %>%
  na.omit() %>%
  FilterSpectra(df = .,
                filter = TRUE,
                return.distances = TRUE,
                num.col.before.spectra = 5,
                window.size = 15)
filtered.test[1:5, c(1:5, (ncol(filtered.test)-5):ncol(filtered.test))]
```

---

FormatCV — *Format multiple trials with or without overlapping genotypes into training and test sets according to user-provided cross validation scheme*

---

#### Description

Standalone function that is also used within TrainSpectralModel to divide trials or studies into training and test sets based on overlap in trial environments and genotype entries

#### Usage

```
FormatCV(trial1, trial2, trial3 = NULL, cv.scheme, seed = NULL,
  remove.genotype = FALSE)
```

#### Arguments

`trial1` : `data.frame` object that is for use only when `cv.scheme` is provided. Contains the trial to be tested in subsequent model training functions. The first column contains unique identifiers, second contains genotypes, third contains reference values, followed by spectral columns. Include no other columns to right of spectra! Column names of spectra must start with "X", reference column must be named "reference", and genotype column must be named "genotype".

`trial2` : `data.frame` object that is for use only when `cv.scheme` is provided. This data.frame contains a trial that has overlapping genotypes with `trial1` but that were grown in a different site/year (different environment). Formatting must be consistent with `trial1`.

`trial3` : `data.frame` object that is for use only when `cv.scheme` is provided. This data.frame contains a trial that may or may not contain genotypes that overlap with `trial1`. Formatting must be consistent with `trial1`.

`cv.scheme` : A cross validation (CV) scheme from Jarquín et al., 2017. Options for cv.scheme include:

- "CV1": untested lines in tested environments
- "CV2": tested lines in tested environments
- "CV0": tested lines in untested environments
- "CV00": untested lines in untested environments

`seed` : Number used in the function `set.seed()` for reproducible randomization. If `NULL`, no seed is set. Default is `NULL`.

`remove.genotype` : boolean that, if `TRUE`, removes the "genotype" column is removed from the output `data.frame`. Default is `FALSE`.

#### Details

Use of a cross-validation scheme requires a column in the input `data.frame` named "genotype" to ensure proper sorting of training and test sets. Variables `trial1` and `trial2` are required, while `trial 3` is optional.

#### Value

List of data.frames (training set, test set) compiled according to user-provided cross validation scheme.

#### Author(s)

Jenna Hershberger <``>

#### Examples

```
# Must have a column called "genotype", so we'll create a fake one for now
# We will use CV00, which does not require any overlap in genotypes
# In real scenarios, CV schemes that rely on genotypes should not be applied when
# genotypes are unknown, as in this case.
library(magrittr)
trials <- ikeogu.2017 %>%
    dplyr::mutate(genotype = 1:nrow(ikeogu.2017)) %>% # fake for this example
    dplyr::rename(reference = DMC.oven) %>%
    dplyr::select(study.name, sample.id, genotype, reference,
                  dplyr::starts_with("X"))
trial1 <- trials %>%
  dplyr::filter(study.name == "C16Mcal") %>%
  dplyr::select(-study.name)
trial2 <- trials %>%
  dplyr::filter(study.name == "C16Mval") %>%
  dplyr::select(-study.name)
cv.list <- FormatCV(trial1 = trial1, trial2 = trial2, cv.scheme = "CV00",
                    remove.genotype = TRUE)
cv.list[[1]][1:5, 1:5]
```

---

`ikeogu.2017` *Example vis-NIRS and reference dataset*

---

#### Description

The ‘ikeogu.2017‘ data set contains raw vis-NIRS scans, total carotenoid content, and cassava root dry matter content (using the oven method) from the 2017 PLOS One paper by Ikeogu et al. This dataset contains a subset of the original scans and reference values from the supplementary files of the paper. ‘ikeogu.2017‘ is a ‘data.frame‘ that contains the following columns:

- study.name = Name of the study as described in Ikeogu et al. (2017).
- sample.id = Unique identifier for each individual root sample
- DMC.oven = Cassava root dry matter content, the percentage of dry weight relative to fresh weight of a sample after oven drying.
- TCC = Total carotenoid content ($\mu g/g$, unknown whether on a fresh or dry weight basis) as measured by high performance liquid chromatography
- X350:X2500 = spectral reflectance measured with the QualitySpec Trek: S-10016 vis-NIR spectrometer. Each cell represents the mean of 150 scans on a single root at a single wavelength.

##### Usage

```
ikeogu.2017
```

##### Format

An object of class `tbl_df` (inherits from `tbl`, `data.frame`) with 175 rows and 2155 columns.

##### Author(s)

Original authors: Ikeogu, U.N., F. Davrieux, D. Dufour, H. Ceballos, C.N. Egesi, and J. Jannink. Reformatted by Jenna Hershberger.

##### Examples

```
library(magrittr)
library(ggplot2)
data(ikeogu.2017)
ikeogu.2017[1:10,1:10]
ikeogu.2017 %>%
    dplyr::select(-starts_with("X")) %>%
    dplyr::group_by(study.name) %>%
    tidyr::gather(trait, value, c(DMC.oven:TCC), na.rm = TRUE) %>%
    ggplot2::ggplot(aes(x = study.name, y = value, fill = study.name)) +
        facet_wrap(~ trait, scales = 'free_y', nrow = 2) +
        geom_boxplot()
```

---

#### PlotSpectra — *Plot spectral data, highlighting outliers as identified using Mahalanobis distance*

---

##### Description

Generates a `ggplot` object of given spectra, with wavelength on the x axis and given spectral values on the y. Mahalanobis distance is used to calculate outliers, which are both identified on the plot. Rows from the original dataframe are printed to the console for each outlier that is identified.

##### Usage

```
PlotSpectra(input.df, wavelengths, num.col.before.spectra = 1,
  window.size = 10, verbose = TRUE)
```

#### Arguments

input.df
: `data.frame` object containing columns of spectra. Spectral columns must be labeled with an "X" and then the wavelength (example: "X740" = 740nm). Left-most column must be unique ID. May also contain columns of metadata between the unique ID and spectral columns. Cannot contain any missing values

wavelengths
: List of wavelengths (numerical format) represented by each spectral column in `input.df`

num.col.before.spectra
: Number of columns to the left of the spectral matrix (including unique ID). Default is 1.

window.size
: number defining the size of window to use when calculating the covariance of the spectra (required to calculate Mahalanobis distance). Default is 10.

verbose
: If `TRUE`, the number of rows removed through filtering will be printed to the console. Default is `TRUE`.

#### Value

If verbose, prints unique ID and metadata for rows identified as outliers. Returns plot of spectral data with non-outliers in blue and outliers in red. X-axis is wavelengths and y-axis is spectral values.

#### Author(s)

Jenna Hershberger <>

#### Examples

```
library(magrittr)
ikeogu.2017 %>%
  dplyr::rename(unique.id = sample.id) %>%
  dplyr::select(unique.id, dplyr::everything(), -TCC) %>%
  na.omit() %>%
  PlotSpectra(input.df = .,
              wavelengths = 350:2500,
              num.col.before.spectra = 5,
              window.size = 15)
```

---

### PredictFromSavedModel *Use provided model object to predict trait values with input dataset*

---

#### Description

Loads an existing model and cross-validation performance statistics (created with `SaveModel`) and makes predictions based on new spectra.

#### Usage

```
PredictFromSavedModel(input.data, model.stats.location, model.location,
  wavelengths = 740:1070, model.method = "pls")
```

#### Arguments

`input.data` data.frame object of spectral data for input into a spectral prediction model. First column contains unique identifiers followed by spectral columns. Include no other columns to right of spectra! Column names of spectra must start with "X".

`model.stats.location` String containing file path (including file name) to save location of "(model.name)_stats.csv" as output from the SaveModel function.

`model.location` String containing file path (including file name) to location where the trained model ("(model.name).Rds") was saved as output by the `SaveModel` function.

`wavelengths` List of wavelengths represented by each column in `input.data`

`model.method` Model type to use for training. Valid options include:

- "pls": Partial least squares regression (Default)
- "rf": Random forest
- "svmLinear": Support vector machine with linear kernel
- "svmRadial": Support vector machine with radial kernel

#### Value

`data.frame` object of predictions for each sample (row). First column is unique identifier supplied by `input.data` and second is predicted values

#### Author(s)

Jenna Hershberger <>

#### Examples

```
## Not run:
ikeogu.2017 %>%
  dplyr::select(sample.id, dplyr::starts_with("X")) %>%
  PredictFromSavedModel(input.data = .,
                        model.stats.location = paste0(getwd(),
                                                      "/my_model_stats.csv"),
                        model.location = paste0(getwd(), "/my_model.Rds"),
                        wavelengths = 350:2500)

## End(Not run)
```

SaveModel *Save spectral prediction model and model performance statistics*

#### Description

Saves spectral prediction model and model statistics to `model.save.folder` as `model.name.Rds` and `model.name_stats.csv` respectively

#### Usage

```
SaveModel(df, save.model = TRUE, autoselect.preprocessing = TRUE,
  preprocessing.method = NULL, model.save.folder = NULL,
  model.name = "PredictionModel", best.model.metric = "RMSE",
  tune.length = 50, model.method = "pls", num.iterations = 10,
  wavelengths = 740:1070, stratified.sampling = TRUE,
  cv.scheme = NULL, trial1 = NULL, trial2 = NULL, trial3 = NULL,
  verbose = TRUE)
```

#### Arguments

`df` — `data.frame` object. First column contains unique identifiers, second contains reference values, followed by spectral columns. Include no other columns to right of spectra! Column names of spectra must start with "X" and reference column must be named "reference"

`save.model` — If `TRUE`, the trained model will be saved in .Rds format to the location specified by `model.save.folder`. If `FALSE`, model will be output by function but will not save to file. Default is `TRUE`.

`autoselect.preprocessing` — Boolean that, if `TRUE`, will choose the preprocessing method for the saved model using the `best.model.metric`. If `FALSE`, the user must supply the preprocessing method (1-12, see `DoPreprocessing()` documentation for more information) of the saved model. Default is `TRUE`.

`preprocessing.method` — Number or list of numbers 1:13 corresponding to desired pretreatment method(s):

- 1 = raw data (default)
- 2 = standard normal variate (SNV)
- 3 = SNV and first derivative
- 4 = SNV and second derivative
- 5 = first derivative
- 6 = second derivative
- 7 = Savitzky–Golay filter (SG)
- 8 = SNV and SG
- 9 = gap segment derivative (window size = 11)
- 10 = SG and first derivative (window size = 5)
- 11 = SG and first derivative (window size = 11)

- 12 = SG and second derivative (window size = 5)
- 13 = SG and second derivative (window size = 11)

`model.save.folder`
: Path to folder where model will be saved. If not provided, will save to working directory.

`model.name`
: Name that model will be saved as in `model.save.folder`. Default is "PredictionModel".

`best.model.metric`
: Metric used to decide which model is best. Must be either "RMSE" or "Rsquared"

`tune.length`
: Number delineating search space for tuning of the PLSR hyperparameter `ncomp`. Default is 50.

`model.method`
: Model type to use for training. Valid options include:
  - "pls": Partial least squares regression (Default)
  - "rf": Random forest
  - "svmLinear": Support vector machine with linear kernel
  - "svmRadial": Support vector machine with radial kernel

`num.iterations`
: Number of training iterations to perform

`wavelengths`
: List of wavelengths represented by each column in `df`

`stratified.sampling`
: If `TRUE`, training and test sets will be selected using stratified random sampling. This term is only used if `test.data == NULL`. Default is `TRUE`.

`cv.scheme`
: A cross validation (CV) scheme from Jarquín et al., 2017. Options for cv.scheme include:
  - "CV1": untested lines in tested environments
  - "CV2": tested lines in tested environments
  - "CV0": tested lines in untested environments
  - "CV00": untested lines in untested environments

`trial1`
: `data.frame` object that is for use only when `cv.scheme` is provided. Contains the trial to be tested in subsequent model training functions. The first column contains unique identifiers, second contains genotypes, third contains reference values, followed by spectral columns. Include no other columns to right of spectra! Column names of spectra must start with "X", reference column must be named "reference", and genotype column must be named "genotype".

`trial2`
: `data.frame` object that is for use only when `cv.scheme` is provided. This data.frame contains a trial that has overlapping genotypes with `trial1` but that were grown in a different site/year (different environment). Formatting must be consistent with `trial1`.

`trial3`
: `data.frame` object that is for use only when `cv.scheme` is provided. This data.frame contains a trial that may or may not contain genotypes that overlap with `trial1`. Formatting must be consistent with `trial1`.

`verbose`
: If `TRUE`, the number of rows removed through filtering will be printed to the console. Default is `TRUE`.

#### Details

Wrapper that uses DoPreprocessing, FormatCV, and TrainSpectralModel functions.

#### Value

List of model stats (in `data.frame`) and trained model object. Saves both to `model.save.folder` as well. To use optimally trained model for predictions, use tuned parameters from `$bestTune`

#### Author(s)

Jenna Hershberger <>

#### Examples

```
library(magrittr)
test.model <- ikeogu.2017 %>%
  dplyr::filter(study.name == "C16Mcal") %>%
  dplyr::rename(reference = DMC.oven) %>%
  dplyr::select(sample.id, reference, dplyr::starts_with("X")) %>%
  na.omit() %>%
  SaveModel(df = ., save.model = FALSE,
            autoselect.preprocessing = TRUE,
            model.name = "my_prediction_model",
            tune.length = 50, num.iterations = 10,
            wavelengths = 350:2500)
summary(test.model[1])
test.model[2]
```

---

### TestModelPerformance *Test the performance of spectral models*

---

#### Description

Wrapper that trains models based spectral data to predict reference values and reports model performance statistics

#### Usage

```
TestModelPerformance(train.data, num.iterations, test.data = NULL,
  preprocessing = TRUE, wavelengths = 740:1070, tune.length = 50,
  model.method = "pls", output.summary = TRUE,
  rf.variable.importance = FALSE, stratified.sampling = TRUE,
  cv.scheme = NULL, trial1 = NULL, trial2 = NULL, trial3 = NULL,
  split.test = FALSE, verbose = TRUE)
```

**Arguments**

`train.data` — `data.frame` object of spectral data for input into a spectral prediction model. First column contains unique identifiers, second contains reference values, followed by spectral columns. Include no other columns to right of spectra! Column names of spectra must start with "X" and reference column must be named "reference".

`num.iterations` — Number of training iterations to perform

`test.data` — `data.frame` with same specifications as `df`. Use if specific test set is desired for hyperparameter tuning. If `NULL`, function will automatically train with a stratified sample of 70%. Default is `NULL`.

`preprocessing` — If `TRUE`, 12 preprocessing methods will be applied and their performance analyzed. If `FALSE`, input data is analyzed as is (raw). Default is `FALSE`.

`wavelengths` — List of wavelengths represented by each column in `train.data`

`tune.length` — Number delineating search space for tuning of the PLSR hyperparameter `ncomp`. Default is 50.

`model.method` — Model type to use for training. Valid options include:

- "pls": Partial least squares regression (Default)
- "rf": Random forest
- "svmLinear": Support vector machine with linear kernel
- "svmRadial": Support vector machine with radial kernel

`output.summary` — boolean that controls function output.

- If `TRUE`, a summary df will be output (1st row = means, 2nd row = standard deviations). Default is `TRUE`.
- If `FALSE`, entire results data frame will be output

`rf.variable.importance` — boolean that:

- If `TRUE`, `model.method` must be set to "rf". Returns a list with a model performance `data.frame` and a second `data.frame` with variable importance values for each wavelength for each training iteration. If `return.model` is also `TRUE`, returns list of three elements with trained model first, model performance second, and variable importance last. Dimensions are `nrow = num.iterations`, `ncol = length(wavelengths)`.
- If `FALSE`, no variable importance is returned. Default is `FALSE`.

`stratified.sampling` — If `TRUE`, training and test sets will be selected using stratified random sampling. This term is only used if `test.data == NULL`. Default is `TRUE`.

`cv.scheme` — A cross validation (CV) scheme from Jarquín et al., 2017. Options for cv.scheme include:

- "CV1": untested lines in tested environments
- "CV2": tested lines in tested environments
- "CV0": tested lines in untested environments
- "CV00": untested lines in untested environments

- `trial1` — `data.frame` object that is for use only when `cv.scheme` is provided. Contains the trial to be tested in subsequent model training functions. The first column contains unique identifiers, second contains genotypes, third contains reference values, followed by spectral columns. Include no other columns to right of spectra! Column names of spectra must start with "X", reference column must be named "reference", and genotype column must be named "genotype".
- `trial2` — `data.frame` object that is for use only when `cv.scheme` is provided. This data.frame contains a trial that has overlapping genotypes with `trial1` but that were grown in a different site/year (different environment). Formatting must be consistent with `trial1`.
- `trial3` — `data.frame` object that is for use only when `cv.scheme` is provided. This data.frame contains a trial that may or may not contain genotypes that overlap with `trial1`. Formatting must be consistent with `trial1`.
- `split.test` — boolean that allows for a fixed training set and a split test set. Example// train model on data from two breeding programs and a stratified subset (70%) of a third and test on the remaining samples (30%) of the third. If `FALSE`, the entire provided test set `test.data` will remain as a testing set or if none is provided, 30% of the provided `train.data` will be used for testing. Default is `FALSE`.
- `verbose` — If `TRUE`, the number of rows removed through filtering will be printed to the console. Default is `TRUE`.

#### Details

Calls `DoPreprocessing`, `FormatCV`, and `TrainSpectralModel` functions.

#### Value

`data.frame` with model performance statistics in summary format (2 rows, one with mean and one with standard deviation of all training iterations) or in long format (number of rows = num.iterations). **Note** if `preprocessing` = `TRUE`, only the first mean of summary statistics for all iterations of training are provided for each technique. Included summary statistics:

- Tuned parameters depending on the model algorithm:
  - **Best.n.comp**, the best number of components
  - **Best.ntree**, the best number of trees in an RF model
  - **Best.mtry**, the best number of variables to include at every decision point in an RF model
- **RMSECV**, the root mean squared error of cross-validation
- **R2cv**, the coefficient of multiple determination of cross-validation for PLSR models
- **RMSEP**, the root mean squared error of prediction
- **R2p**, the squared Pearson's correlation between predicted and observed test set values
- **RPD**, the ratio of standard deviation of observed test set values to RMSEP
- **RPIQ**, the ratio of performance to interquartile difference
- **CCC**, the concordance correlation coefficient
- **Bias**, the average difference between the predicted and observed values
- **SEP**, the standard error of prediction
- **R2sp**, the squared Spearman's rank correlation between predicted and observed test set values

#### Author(s)

Jenna Hershberger <>

#### Examples

```
library(magrittr)
ikeogu.2017 %>%
  dplyr::rename(reference = DMC.oven) %>%
  dplyr::rename(unique.id = sample.id) %>%
  dplyr::select(unique.id, reference, dplyr::starts_with("X")) %>%
  na.omit() %>%
  TestModelPerformance(train.data = .,
                       tune.length = 3,
                       num.iterations = 3,
                       preprocessing = FALSE,
                       wavelengths = 350:2500)
```

### Index

∗ **datasets**
    ikeogu.2017, 7

AggregateSpectra, 2

DoPreprocessing, 3, *11*, *13*, *15*

FilterSpectra, *2*, 4
FormatCV, 5, *13*, *15*

ggplot, *8*

ikeogu.2017, 7

PlotSpectra, 8
PredictFromSavedModel, 9

SaveModel, *9*, *10*, 11

TestModelPerformance, 13
TrainSpectralModel, *5*, *13*, *15*
